# Detection and quantification of planar traveling waves in the EEG using spherical phase fitting

**DOI:** 10.64898/2025.12.03.692197

**Authors:** Jakob C. B. Schwenk, Andrea Alamia

**Author notes:** Corresponding author : Jakob Schwenk.

## Abstract

Recent years have seen an increasing interest in the spatiotemporal dynamics of oscillatory brain activity. Recordings at various scales have shown oscillations propagating as waves over cortical space, both at small and large scales. The most common waves observed in the EEG are planar waves, formed by a synchronized wavefront propagating in a consistent direction (e.g., posterior-anterior). These waves have been linked to diverse perceptual and cognitive measures. However, their quantification has faced issues due to the inherently noisy EEG signal and its ambiguous spatial localization. Existing methods employ restrictive windows of analysis (e.g., lines of electrodes) or rely on wave motifs extracted from the data. A specific algorithm for detecting planar waves is still lacking. Here, we present a comprehensive analysis pipeline for this purpose, comprising three steps: first, genuine oscillatory activity is extracted from the EEG as clusters. The spatial phase gradient is then fit using a spherical wave model. Finally, stable waves are extracted from the time series of fits. We validate our method using simulations over a physiological range of parameters. Using a forward model, we test EEG wave detection for propagation along different pathways at the source level. Lastly, we apply our analysis to real EEG recordings, targeting alpha oscillations during visual stimulation and at rest. In summary, our method provides a reliable algorithm for detecting planar waves in the EEG. Given the emerging functional roles of traveling waves in perception and cognition, this could potentially be utilized in a wide range of future studies.

## Introduction

The spatiotemporal dynamics of oscillatory brain activity have gained substantial interest in recent years. A prominent phenomenon is that of traveling waves, i.e., the synchronized propagation of neural activity over cortical space (see (Muller et al., 2018) for an overview). Studies using intracortical and cortical surface recordings have reported a wide array of traveling wave phenomena, both at meso-scale (within areas) and as large-scale waves spanning multiple areas or entire hemispheres. Broadly, these can be grouped into waves occurring spontaneously (Bahramisharif et al., 2013; Davis et al., 2020; Halgren et al., 2019; Zhang et al., 2018), and those induced by sensory input (Chemla et al., 2019; Muller et al., 2014, 2018; Sato et al., 2012) or related to the execution of movements (Balasubramanian et al., 2020; Best et al., 2017; Rubino et al., 2006; Takahashi et al., 2011, 2015). The most common patterns observed are synchronized wavefronts traveling in a single direction (planar waves), or expanding out from a single source (radial waves) (for a description of more complex patterns, see, e.g., (Townsend et al., 2015; Townsend & Gong, 2018)).

The question of whether and how exactly cortical waves project to the EEG is not straightforward (as will be discussed further below). However, phenomenologically, waves traveling along the scalp have been observed in a wide array of experimental contexts. Most prominently, these include long-range planar waves in the alpha range (7-13 Hz) whose directional dynamics are modulated by low-level features of visual stimuli (Alamia & VanRullen, 2019; Lozano-Soldevilla & VanRullen, 2019; Pang (庞兆阳) et al., 2020; Schwenk & Alamia, 2025) and actively in relation to cognitive/perceptual tasks (Alamia et al., 2023; Fellinger et al., 2012; Patten et al., 2012; Zeng et al., 2024), with possible correlations to clinical phenomenology (Alamia et al., 2024). A recent line of investigations has also shown that rhythmically induced meso-scale waves traveling over retinotopic space in early visual areas can elicit global wave patterns in the EEG (Fakche & Dugué, 2024; Grabot et al., 2025; Petras et al., 2025).

This ubiquity of traveling waves in the EEG highlights the need for reliable and standardized methods for their quantification. The main challenges faced by this are the inherently noisy nature of the EEG signal, as well as the highly variable occurrence of waves, which can form as transient events of a few cycles (see, e.g., (Patten et al., 2012)) up to sustained oscillations in the range of multiple seconds (e.g. in (Pang (庞兆阳) et al., 2020)). Existing methods can be broadly categorized into three categories. The first of these uses the two-dimensional Fast-Fourier-Transform (2D-FFT; first adapted for wave quantification by (Alamia & VanRullen, 2019)) to characterize planar waves along a pre-defined line of electrodes. In that approach, waves are quantified as a continuous measure, derived from the spectral power associated with either direction on the axis of propagation (e.g., anterior-posterior on the midline). While easy to implement, this method remains restricted to a single dimension in sensor space, and offers low temporal resolution. Moreover, the continuous measure of wave power, although powerful in aggregate comparisons (e.g., between conditions), is not well suited for detecting individual wave events at the level of single trials. The second category of methods uses linear fits on the map of oscillatory phase over sensors. Most studies that have employed this approach have used simple line fits on selected electrode lines, analogous to the 2D-FFT (Burkitt et al., 2000; Ito et al., 2005; Patten et al., 2012). We have previously extended this to two dimensions, approximating a small region of the scalp by a plane (Schwenk & Alamia, 2025). The new method presented here represents a significant improvement and generalization of this approach.

Lastly, the third category of methods comprises data-driven approaches, wherein dominant spatiotemporal patterns are extracted from the data using decomposition of the phase structure (Alexander et al., 2006, 2013; Li et al., 2025; Patten et al., 2012; Petras et al., 2025). The benefit of these methods is that they are hypothesis-free (as compared to the 2D-FFT or linear fits). However, they are not ideally suited for detecting specific (and potentially rare) wave patterns, as they need to explain a sufficient amount of variance in the data and must be matched with the target wave shape post-hoc. Additionally, waves of the same type may occur transiently at different topographical locations.

Our goal was to develop an analysis algorithm for detecting planar traveling waves in the EEG. Specifically, to improve upon previous model-based approaches (2D-FFT and linear fits), our target criteria were higher resolution in both the temporal and spatial domains, without imposing prior assumptions about the localization of the wave on the scalp. Our proposed algorithm combines oscillatory clustering with an improved wave fitting, which accounts for the spherical shape of the scalp to model planar waves. Its target use case is the analysis of EEG data with clear oscillatory components that surpass background noise and aperiodic activity at a sufficient SNR.

### General description of the method

In this section, we describe our proposed analysis pipeline for detecting and quantifying planar traveling waves. Figure 1 provides a comprehensive overview of all steps, using an example traveling wave event from real EEG data. Broadly, the procedure consists of three steps. The first step is the extraction of clusters of oscillatory activity in the EEG (Figure 1A-C). While traveling waves themselves are defined in phase, we consider (in the absence of *a priori* assumptions) only those waves that are forming from true oscillatory activity, i.e., with power surpassing the aperiodic (1/f) activity. In the second step, the spatial patterns of the phase inside each cluster are fit using a spherical planar wave model (D). Lastly, from the time-course of the best fits, time periods in which the wave parameters are stable are extracted as traveling wave *epochs* (E). Further analysis and hypothesis testing is then more easily performed on these epochs (F), which can be simplified to have scalar values for each wave parameter (e.g., direction, spatial frequency). In the following sections, we will provide a detailed description of each step.

**Figure 1.**
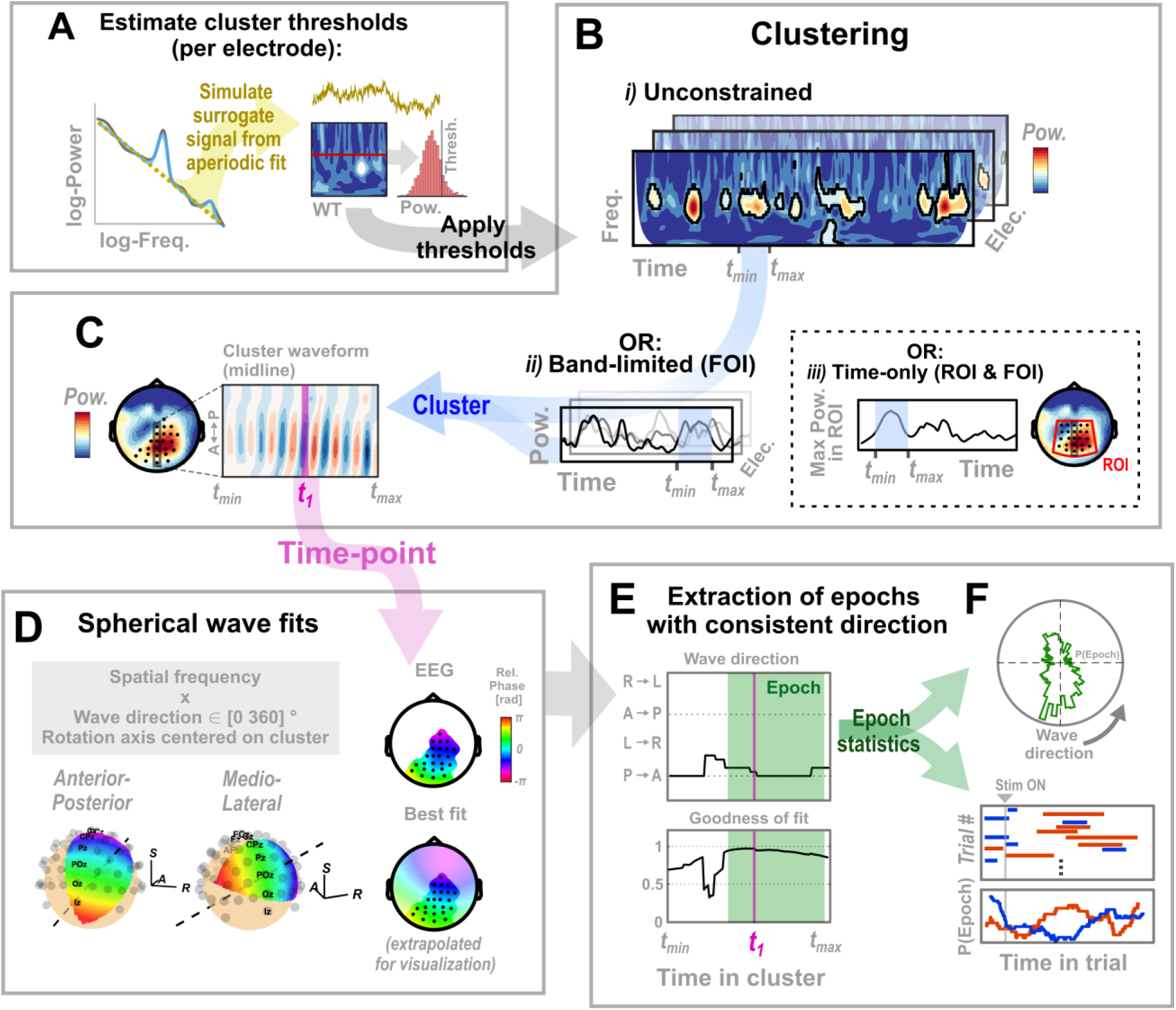
Overview of the proposed algorithm for the detection of planar traveling waves in the EEG. For details on each step, see main text. A: Estimation of power thresholds for the clustering. For each electrode, an aperiodic fit is obtained (Donoghue et al., 2020) to simulate surrogate data with chance-level oscillatory power. The wavelet transform (WT) of this surrogate data is used to estimate a random distribution of power expected at each frequency. To this, a percentile threshold is applied to yield consecutive regions in time x frequency x electrode-space that show true oscillatory activity. B: Extraction of oscillatory clusters. Clusters are defined as contiguous regions in the full WT of the EEG that surpass the thresholds estimated in A. These can be defined either in all three dimensions (i) or limited in frequency (ii) or frequency & electrode-space (iii) to predefined regions of interest. In cases ii) and iii), the measure for power inside the FOI/ROI is collapsed to a single value (e.g., mean over FOI, maximum within ROI), both for the surrogate and the real data. C: The spatiotemporal pattern of activity inside each cluster is then used to evaluate the presence of traveling waves. The right panel shows the topographical distribution of power for an example cluster defined over posterior sensors (highlighted). The left panel shows the activity for a line of electrodes inside the cluster over time. A forward wave pattern is already visible. D: For each time-point inside the cluster, the spatial pattern of phases is compared with spherical wave templates, each defined by a wave direction and a spatial frequency (for details, see main text). The left panels display templates for an anterior-posterior (forward) and a medio-lateral (left-to-right) wave as examples, for the same cluster shown in C. The axis on which the line of propagation is rotated to vary wave direction is defined by the center of the cluster (dashed line). All templates are limited to a central +/-45° perpendicular to the wave direction to avoid the singularities at the poles. The right panels show the spatial phase maps for the data (top) and the corresponding selected best fit (bottom), extrapolated to the full head for better visualization of the wave direction. Phases are all relative to the mean across the cluster region. Axes legend: S – Superior, A – Anterior, R – Right. E: From the time-course of best fits within the cluster, waves are finally extracted as ‘epochs’, defined as time-periods in which the wave direction and goodness of fit remain sufficiently stable. Panels show the time-courses for these parameters for the example cluster, with an extracted epoch in the shaded green area. Wave directions: L/R – left/right, A/P – anterior/posterior. F: Statistics of these epochs (with scalar values for the wave parameters) can be used for hypothesis testing and exploration, for example by analyzing frequency distributions of wave direction (top) or comparing probabilities of different wave types (e.g., FW/BW) occurring throughout a trial.

### Extraction of oscillatory clusters

We base our detection of true oscillatory activity in the EEG on the spectral parametrization procedure introduced by Donoghue and colleagues (Donoghue et al., 2020). This algorithm models the observed power spectral density (PSD) of the EEG signal as the sum of true oscillatory peaks (at multiple frequencies) and an aperiodic component. This aperiodic component is estimated by a log-linear fit to the PSD, and subtracted from the full spectrum to yield the true oscillatory activity attributable to neural sources. For our purpose, we are interested in the aperiodic activity to get an estimate of the power at each frequency that can be considered chance-level. Specifically, to extract time periods of oscillatory activity at high temporal resolution, we base our full analysis on the continuous wavelet transform (WT). Since the aperiodic fit is based on a stationary measure of the PSD that does not flexibly translate into power in the WT, we use simulated data to obtain our estimate of the chance-level distribution. This is illustrated in Figure 1A.

First, for each electrode, the PSD is obtained using a standard time-frequency decomposition (Fast Fourier Transform (FFT) with the multi-taper method) based on segmenting the data into short segments. For the results presented here, we used segments of 2 seconds with an overlap of 0.5 seconds. From this, the aperiodic component is estimated as described in the original publication (Donoghue et al., 2020). Here, we utilize the implementation provided by the fieldtrip toolbox (Oostenveld et al., 2011), applied to the full original dataset. Depending on the research question and level of noise in the target data, the fit can also be obtained from a separate baseline condition (e.g., resting state). Using an inverse FFT, we then simulate surrogate chance-level data (90 secs, for each electrode) with the estimated power slope and random phase values, and finally compute the wavelet transform (WT), matching the analysis of the real data. The distribution of wavelet-power at each frequency serves as the basis in the following extraction of oscillatory clusters.

The rationale behind our cluster-based approach is that only oscillations with sufficient power can be assumed to stem from neural sources. However, for a given oscillatory event, neighboring points in time or sensor-space that don’t surpass the initial threshold are likely still interpretable. Therefore, we adopt a dual thresholding procedure. In the first step, candidate clusters are detected as contiguous time-frequency points where the wavelet power surpasses an initial high threshold (*thresh_high_,* set at 95% of the random distribution for the main results presented here). Each cluster is then extended to the time period and sensor region within which values surpass a lower threshold (*thresh_low_*, set at 70% in the following). Importantly, the limits in the frequency domain are retained from the higher thresholding, to avoid broadband clusters comprising multiple oscillatory peaks. Note that the extension to the lower-threshold bounds also leads to clusters being merged that were separated (in either dimension) in the initial clustering.

In an optional additional step, clusters that are distinct (at both thresholds) may be merged if they are separated primarily in electrode space. This is achieved by applying a threshold to the amount of overlap in time and frequency-space between all pairs of clusters. This step enables the classification of oscillatory clusters with two (or more) peaks as a wave, for example, as generated by two oscillating dipoles that are sufficiently separated in source space. The limitations of our wave quantification for these types of spatiotemporal patterns are discussed below (see *Results: Source-level wave simulations with oscillating dipoles*).

To filter out transient high-amplitude, non-oscillatory events, clusters are excluded from subsequent fitting if their extent in the frequency domain is very large (based on the log_10_-ratio of the frequency boundaries; threshold of log_10_(f_max_/f_min_) > 0.5 used here).

Lastly, given that our fit requires sufficient spatial coverage, we also exclude all clusters that are too small in electrode space from further analysis (the minimum cluster size used here is 4 electrodes; higher-density recordings may require larger values).

### Alternative lower-dimensional cluster-extraction (FOI and/or ROI)

If the waves of interest are in a specific (narrow) frequency band, the cluster extraction can be adapted to a frequency region-of-interest (FOI, e.g., alpha: 7-13 Hz), illustrated in Figure 1B, ii). For this, the wavelet power at each step is averaged across the FOI, resulting in two-dimensional data (electrode x time). Note that the aperiodic fit still needs to be performed as before on the full spectrum to obtain a reliable estimate of the chance-level distribution of the FOI-averaged power. This procedure increases sensitivity to include oscillatory activity in that band. However, as frequency information is lost, the resulting clusters can be influenced by activity at neighboring frequency bands. For this reason, FOI-based clustering should only be applied with a fixed hypothesis about the chosen frequency band. Depending on the research question, this could be, for example, the individual alpha frequency (IAF) or specific oscillatory peaks detected by the spectral parametrization (Donoghue et al., 2020).

Following the same logic, the analysis can be further restricted to an electrode region-of-interest (ROI), illustrated in Figure 1B, iii). Here, the *a priori* assumption would be that waves occur always in the same frequency range and in the same region of the scalp (but, e.g., changing wave direction across conditions). In this case, clustering is performed in time only (1D) after first averaging over the FOI and then pooling over the ROI (e.g., taking the maximum). Note that, unlike the FOI-only (2D) clustering, this method predefines a constant set of electrodes to be used for the fit (this case is not included in the illustration in Figure 1D).

### Definition of the phase gradient for spherical planar waves

Following the extraction of oscillatory clusters, the signal inside each cluster is fit with a spherical phase gradient model to determine whether its spatiotemporal distribution is consistent with a traveling wave. Our fitting procedure is based on the planar traveling wave detection mehod developed by Zhang et al. for ECoG recordings (Zhang et al., 2018). However, we adapt the underlying phase gradient model to the spherical surface of the scalp, as follows.

First, the sensor positions in 3-dimensional space are shifted to the nearest points on a common sphere, based on a least-squares fit of all sensors. For the results presented here, we used the standard 10/5 arrangement provided by the fieldtrip toolbox (mean absolute residual of the sphere approximation of 5% of the sphere radius across all 97 electrodes in the set).

We define the phase gradient along the continuous angular elevation spanned by two of the spherical axes. In other words, our template assumes that each wave travels along some *great circle* on the sphere (an example of this being the earth’s equator line), and along smaller circles running in parallel to it (up to some minimal distance from the singularity at the poles, which are masked out, as detailed below).

The formulation of the phase gradient as a function of θ and *ξ* is intuitively best understood as a sequence of 3 rotations: the first two align the axis of rotation with the center of the cluster, while the third rotates on that adjusted axis to the desired wave direction.

The Euler-/Rodrigues-rotation for a vector *p* of positions in 3D space is defined as:

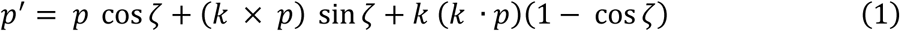

where *p’* is the vector of rotated positions, is the angle of rotation and k is a unit vector describing the axis of rotation.

We follow the convention that the sensor positions are aligned in 3D space such that the first two dimensions extend, respectively, mediolaterally (M-L, *x*) and anterior to posterior (A-P, *y*), while the third dimension extends from inferior to superior (I-S, *z*). The three rotational angles, following common terminology for head rotations, are yaw (in the x-y plane), pitch (y-z), and roll (x-z).

From the sensor positions *p:*

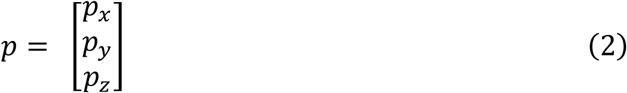

we obtain the cluster-centered positions *q* by rotating first on the *y-* and then on the (rotated) x-axis (adjusting for pitch and roll). For each cluster, the values of these rotations are chosen to align the z-axis with the topographical region of highest oscillatory power. Specifically, we use the pitch and roll of the topographical center of mass (power-weighted average position) of the top 10% of power values inside the cluster topography. This allows the rotational axis to be centered between potential separate peaks, for example, in the case of a bimodal distribution. To avoid ambiguity in the interpretation of wave direction, we limit the axial adjustment in both pitch and roll to values in the range +/-45 degrees (i.e., covering only the top half of the head-sphere).

The resulting vector *q* is now aligned such that the phase template of a wave with direction θ is a linear scaling of the pitch (y-z inclination) of the rotation of *q* by θ around the z-axis:

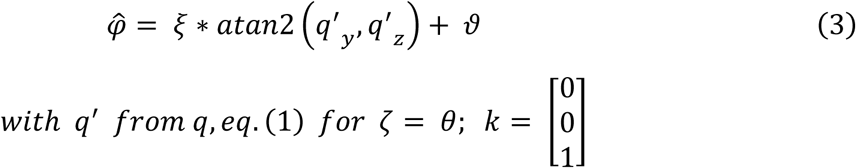

where *ϕ^* is the vector of predicted phases at the original set of sensor positions in *p,* and ϑ is a scalar phase offset. The left panels in Figure 1D illustrate the phase templates for two waves with orthogonal wave directions.

### Fitting procedure

The fit is applied to the observed phase gradients within the limits (in time/frequency/sensor-space) of each cluster separately. For this, the time-domain signal corresponding to the cluster is recovered using the inverse wavelet-transform (after setting all values outside the cluster to zero). The instantaneous phase is then extracted from the Hilbert transform of the filtered signal and re-referenced to the mean phase angle for each timepoint.

For each time-point inside the cluster (and for all electrodes active at that time-point), the best fit is then evaluated iteratively from the 2D space over all combinations of wave direction θ ∈ [0 2π] and spatial frequency *ξ* (which can be bounded to some biologically plausible range of propagation velocities).

The best fit is determined by maximizing the vector length of the summed residuals, given by:

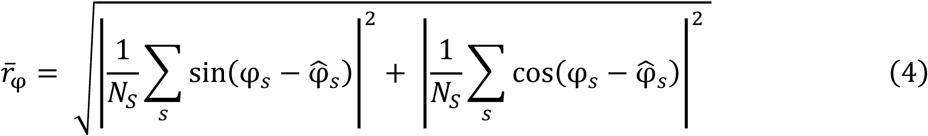

where *φ* and *ϕ^* are the observed and predicted phase values, respectively, subscript *s* indexes sensors and *N_S_* is the number of sensors active in the cluster (at the given timepoint).

We use the adjusted circular correlation ρ between the predicted and observed phases as our measure of goodness of fit. Following Zhang et al. (Zhang et al., 2018), we denote this as the phase gradient directionality index (PGD):

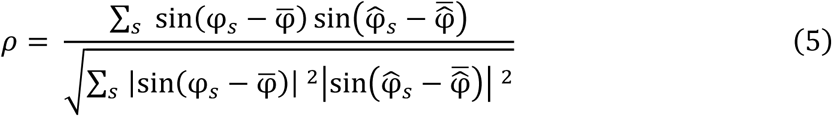

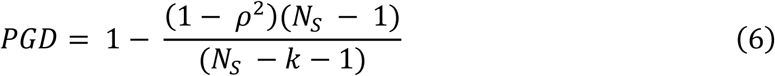

with notation following eq. (4), φ̅, φ^̅ denoting the average between sensors of observed and predicted phases, respectively, and *k = 3* given by the number of independent regressors.

The set of available electrodes used for the fit is defined by the cluster (note that this can vary between time-points). The set is further restricted for each fit by the wave template, which we limit to an angle of +/-45° on the axis orthogonal to the wave direction to avoid phase singularities at the poles and large distortions in propagation velocity (the resulting shape of the template can be seen in Figure 1D, left panels). To avoid unstable fits resulting from low spatial coverage, we invalidate (by setting PGD to a low value) all fits for which the set of included electrodes does not cover a minimum distance in relative phase across the template (set here at 45° or 0.785 rad). This also prevents standing waves (with uniform phase) from being erroneously picked up by fits with very low spatial frequency.

The result of the fitting procedure is a single set of wave parameters (direction θ and spatial frequency *ξ*) and goodness of fit (PGD) for each time point in the cluster (illustrated in Figure 1E).

### Extraction of wave epochs

The time series of best fits for each cluster still contain periods where either no wave was present (low goodness of fit) or wave parameters were inconsistent over time. Therefore, we perform a second-stage clustering, this time on the extracted wave direction and PGD, to yield the final time windows of stable traveling waves, which we denote as *epochs*.

For a given cluster of duration *T_max_*, we iterate through all possible epoch durations *T =* [*T_max,_ T_max_ -1, …, T_min_*], where *T_min_* is the minimum epoch duration considered a stable wave. For the results presented here, we set *T_min_* at 2 oscillatory cycles of the peak frequency of the cluster. An epoch is detected wherever the mean angle of the wave direction within *T* consecutive time-points is maximal and exceeds a given threshold (minimum circular coherence, *CC_min_ = 0.8* used here). Similarly, PGD values must not fall below a certain threshold (here, *PGD_min_ = 0.5*), after interpolating short periods of poor fits (max. consecutive 150 ms, accounting for no more than 30% of the epoch duration). More than one epoch can be detected for a given duration. If no valid epoch is detected, the algorithm is repeated on the next shorter duration.

Finally, for each epoch, we retain all target parameters (mean wave direction, spatial frequency, PGD, peak temporal frequency and power) as scalar values for further analysis.

## Validation methods

In the following sections, we will describe the methods used in this paper to test and validate our proposed algorithm, using simulated and real data.

### Simulation of EEG data

We base our validation of the wave detection algorithm on simulated EEG data. Specifically, we used different configurations of waves (or wave-like activity) simulated at both sensor and source levels, with distinct aims.

### Sensor-level waves

Our sensor-level simulation of waves was designed to test how robustly our algorithm can detect idealized waves in the presence of noise of different origins. For this, we combined three signals: source-level aperiodic activity and sensor-level white noise (at variable mixing proportions), and the sensor-level wave given by the spherical phase gradient defined above.

The source-level noise consisted of five different dipoles, each following independent random sequences with an aperiodic power spectrum (slope 1/f). These sources were projected to the sensor level using a standard forward projection (fieldtrip function *ft_dipolesimulation*) with radially oriented dipoles. Source positions were drawn randomly for each trial from a grid of 20 mm resolution.

Channel-level noise was independently added to the signal at each sensor as white noise.

Lastly, the wave component was added as a pure sine wave (*F = 10 Hz* for all reported simulations) with a constant phase gradient. To simulate a wave localized over occipital areas, wave topography was defined as a von-Mises distribution, with signal amplitude decreasing as a function of angular distance from electrode POz (spread parameter k = 3). Peak signal-to-noise ratios (SNR) for the reported results refer to the ratio of the root-mean-square (RMS) value between the wave component at the peak electrode (POz) and the noise-signal (mean across all sensors).

For a given set of noise parameters, we simulated 50 trials of 3 seconds, with the wave occurring transiently for 1 second (onset t = 1 sec, no ramp-in). Across trials, wave direction was varied randomly. We evaluate performance using the probability (across trials) of the wave being detected as an epoch, and the accuracy of the wave direction estimate. The latter is computed from the absolute residuals in circular space as:

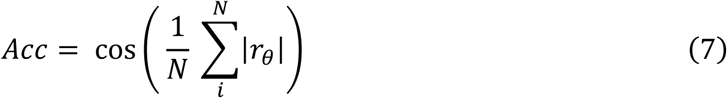

where subscript *i* indexes the trials and *r*_θ_ is the phase difference between the estimated and true wave directions, wrapped to the range [-π, π]. This measure takes values between -1 and 1, where 0 represents a chance-level estimate, 1 represents a perfect estimate, and -1 represents a consistent estimate of the opposite direction.

### Source-level wave simulations with oscillating dipoles

In a second set of simulations, we tested various source-level configurations that could result in planar traveling wave patterns at the sensor-level. Since the true neural substrates underlying traveling waves in the EEG are not fully understood, these simulations are exploratory and not designed to test performance against a ground truth. We follow previous studies, which show that few oscillating sources can generate wave-like patterns if their phase has a consistent offset (Schwenk & Alamia, 2025; Zhigalov & Jensen, 2023).

The simulations employed the same noise background described above, with a fixed mixing proportion of 0.15 (RMS ratio of channel-to-source-level noise). The oscillating (wave-) sources were positioned in different configurations between 7 possible positions (coordinates listed below), which we denote as *pathways*. Each pathway simulates sinusoidal activation propagating at a constant speed along a single stream of areas, following either the dorsal or ventral visual hierarchy. The activity for a given dipole *N* and a wave traveling forward (FW) or backward (BW) is given by:

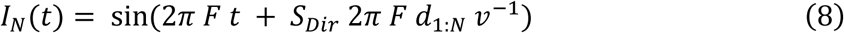

Here, *N* indexes sources, *t* is time, *F* is the temporal frequency of the wave (*F = 10 Hz* for all simulations reported), *v* is the velocity of the wave, *d*_1:*N*_ is the cumulative Euclidian distance between sources from *1* to *N,* and the sign of the phase offset *S*_*Dir*_ ∈ {−1 1} determines the direction of the wave (forward (FW) or backward (BW)).

The dipoles were projected to the EEG in the same manner as the source-level noise. As before, the peak SNR refers to the ratio of the RMS at the sensor with the maximal signal strength. Similarly, we evaluated the results of our wave-detection algorithm again over 50 trials for each set of parameters tested. Between trials, wave direction was split equally between FW and BW. For each trial, the detected wave epoch was classified as FW/BW if the wave direction was within 45° of the respective reference direction. For all pathways between occipital, parietal and frontal sources, these were aligned with the anterior-posterior axis (*θ_ref_ = 0 rad* if FW, *π rad* if BW). For the occipital-temporal pathway, this axis was rotated by 45° towards the active hemisphere. Accuracy of the wave detection was defined as the percentage (across trials) of correctly classified epochs.

The positions for the dipoles were selected based on the AAL atlas (Tzourio-Mazoyer et al., 2002) to simulate activity in occipital, parietal, frontal, and temporal regions. For most pathways, we used a bilateral configuration of sources, with two non-interconnected streams of identical activity in the two hemispheres. The MNI coordinates for these positions are listed in Table 1.

**Table 1.**
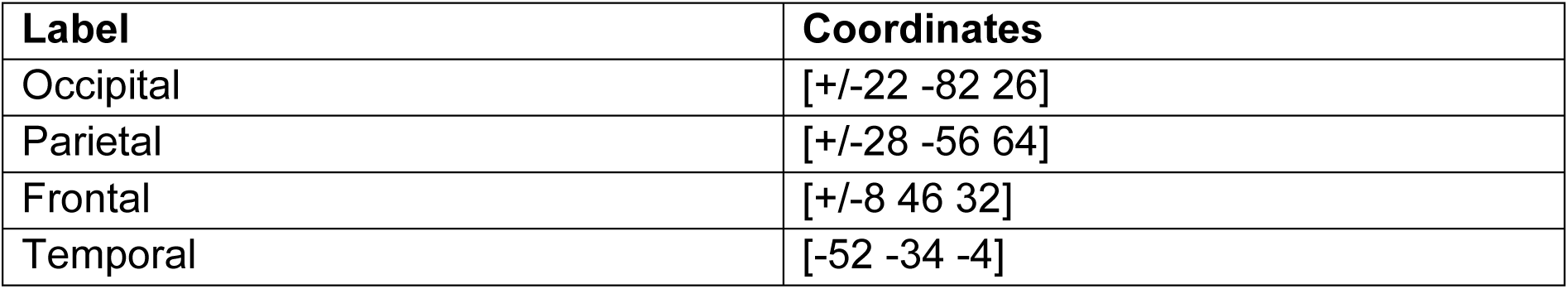
MNI coordinates of dipole positions used for the source-level wave simulations. For each area/label except the temporal dipole, configurations were tested with one source in each hemisphere, therefore x-coordinates are given bilaterally.

| Label | Coordinates |
| --- | --- |
| Occipital | [+/-22 -82 26] |
| Parietal | [+/-28 -56 64] |
| Frontal | [+/-8 46 32] |
| Temporal | [-52 -34 -4] |

### Analysis of real EEG datasets

In addition to our simulation tests, we also applied our algorithm to real human EEG data. Specifically, we first re-analyzed a dataset for which traveling wave dynamics (in the alpha band) have already been extensively studied (Pang (庞兆阳) et al., 2020; re-analyzed using a different method in Schwenk & Alamia, 2025). This served as a confirmatory investigation to further validate our new algorithm. We then also analyzed two publicly available resting-state datasets to test generalizability of our results.

#### Visual stimulation vs. rest: Pang (庞兆阳) et al. (2020)

The experimental procedure underlying this dataset is described in detail in the original publication. In brief, human subjects (N = 13 in the final sample, 6 females) viewed a randomly flickering disk at uniform greyscale luminance presented in the visual periphery (‘Dynamic’ condition in the original paper, comprising 150 trials). The subjects’ task was to covertly monitor the disk for the random appearance of a target stimulus. The visual stimulation lasted 5 seconds on each trial and was followed by another 5 seconds of blank screen. Throughout the entire trial, subjects were instructed to maintain central fixation. All subjects gave written informed consent, and the experimental protocol was approved by the ‘Comité de protection des Personnes Sud Méditerranée 1’ (approval number N°2016-A01937-44).

Signals were recorded using a 64-channel EEG system (BioSemi) at a sampling rate of 1024 Hz and down-sampled to 160 Hz offline. Re-referencing to the average was performed, with high-pass (>1Hz) and band-stop (50 Hz) filters applied. Manual artifact detection and rejection were performed after epoching to the trial time, yielding an average of 135 trials per subject used for the final analyses.

We re-analyzed the data using our new algorithm, which employs the full clustering approach (as described above), for frequencies between 2 and 20 Hz, based on the original 10-second trials (each comprising one consecutive stimulation and one rest period). The spectral parametrization was performed on a re-segmented version of the dataset, using segments of 2 seconds with an overlap of 0.5 seconds.

##### Resting state: Chenot et al. (2024)

This dataset is part of a pre-registered study on hypothesized links between resting-state dynamics and cognitive executive functions. Our analyses in this work are not related to that study. The experimental procedure is described in detail in the published first report (Chenot, Hamery, Truninger, Langer, et al., 2024). The subjects (N = 140 in the final sample, comprising 66 females, with a mean age of 24.72 years) participated in two sessions, the first of which included the EEG resting-state measurements used in this study. For this, subjects were seated in front of a computer screen and instructed to relax, keeping their eyes open or closed in alternating periods of 30 seconds each, for a total of 5 minutes.

EEG signals were recorded on a 64-channel EEG system (BioSemi), at a sampling rate of 512 Hz.

For our analysis, we pre-processed the data as follows. First, an independent component analysis (ICA) was performed (using the EEGLAB *runica* algorithm, (Delorme & Makeig, 2004)), and all components classified as eye-movement-, heart- or muscle-related removed. The cleaned data was then re-referenced to the channel-average, and high-pass (> 1Hz) and band-stop (50 Hz) filters applied. The wave detection algorithm was run using band-filtered (two-dimensional) clustering (FOI: 7-13 Hz) on 5-second, non-overlapping segments of the data. Spectral parametrization was performed on the same data, segmenting it into 2-second intervals with a 0.5-second overlap.

##### Resting state: Babayan et al. (2019)

This EEG dataset is derived from a large, multimodal data repository (MPI Leipzig Mind-Brain-Body database). A comprehensive description of the sample is provided in the original publication. We used only the resting-state EEG measurements for our analyses. These were acquired in a subset of the larger sample from the full study (N = 202 in the final sample used here, comprising 74 females, with a mean age in the 35–40 years range). For each subject, EEG was collected over a 16-minute recording, using a BrainAmp MR plus amplifier with 61 scalp electrodes (Brain Products, Gilching, Germany), sampled at 2.5kHz. The recording sequence consisted of an alternation of 16 blocks, each 60 seconds long, with blocks alternating between eyes-open (EO) and eyes-closed (EC) conditions. Subjects fixated a black cross on a white background on a computer screen during the EO condition.

We used the pre-processed data provided in the data repository for our analyses. For this the data were first down-sampled to 250Hz, and a band-pass filter applied between 1 and 45 Hz. Initial rejection of large artifacts was performed visually. Subsequently, a principal component analysis (PCA) was used to reduce dimensionality of the data to those components explaining 95% of the variance. Lastly, an ICA was performed and artifactual components removed. For more details on the pre-processing, see the original publication (Babayan et al., 2019).

We applied our algorithm using the same settings as for the Chenot et al. dataset (see above).

## Results

The analyses we performed to evaluate the validity of our method consisted of three steps, as detailed below. In the first two sections, we present tests of the algorithm’s performance in controlled simulations of noise and waves defined in sensor space. We then investigate how reliable wave patterns can be detected at the scalp when generated by different source-level configurations. Lastly, we apply our method to one previously analyzed and two publicly available EEG datasets, replicating previous findings on alpha wave direction at rest and during stimulation.

### Estimate of chance-level classification probability

In a first step, we estimated the specificity of our detection algorithm, i.e., the rate of detected wave epochs when evaluated on random noise. For this, we projected aperiodic activity at the source level to a simulated EEG (Figure 2A, for details see Methods, *Simulation of EEG data*). The spatial mixing of these noise signals leads to wider regions of sensors being synchronized in phase, and transient wave-like patterns can form if two or more noise sources have a consistent phase offset over a certain time period. This simulates the physiological noise typically present in real EEGs.

**Figure 2.**
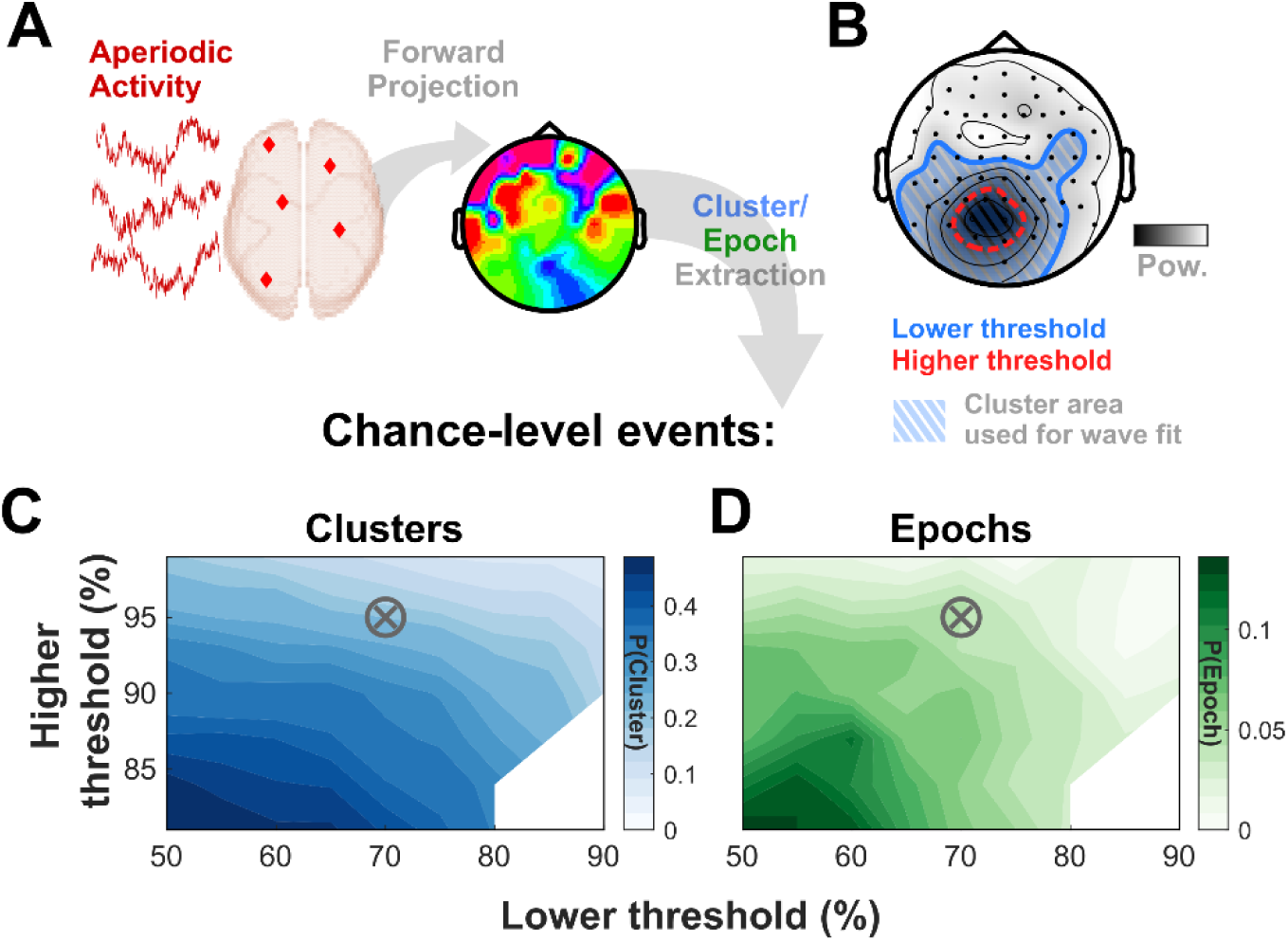
Chance-level probabilities for cluster and epoch occurrence as a function of the two thresholds applied in the cluster detection. A: Procedure used for the simulation of physiological noise. Randomly positioned sources of aperiodic activity were forward-projected to the EEG and clusters/epochs detected on the simulated signal. B: Schematic of the dual-thresholding procedure for cluster-extraction. Wave fits are evaluated in the time and electrode-region defined by the lower threshold (shaded blue area). C, D: Chance-level event rate for clusters and epochs. Probabilities are evaluated as the total proportion of time-points marked as either cluster or epoch. The combination of threshold values used for all following results is marked by a circled cross (70% / 95%).

Our extraction of oscillatory clusters (on which the wave fit is evaluated) is based on a dual thresholding (Figure 2B, for details see Methods, *Extraction of oscillatory clusters*): the first, higher threshold is used for inclusion of the cluster and defines its extent in the frequency domain. The cluster is then extended in electrode-space and time up to a second, lower threshold. Both thresholds are defined as percentiles of the estimated random distribution of oscillatory power at each electrode. Note that the percentiles of those distributions do not correspond to the chance-level percentage of clusters, as detection of the latter takes into account the full connectivity in electrode / time / frequency-space.

We tested our algorithm on simulated noise for a range of values of both thresholds and extracted the probabilities of clusters and epochs (across a set of trials) for each combination. The results are shown in Figure 2.

As expected, the rate of chance-level events decreases with increasing thresholds. Approximately one-fifth of the total time included in clusters was further classified as an epoch (mean ratio P(Epoch)/P(Cluster) = 0.22 for all combinations where either probability was > 0.05). For all results presented below, we chose values of 70% and 95% for the two thresholds, respectively. This choice yields an approximate chance-level detection of wave epochs of 5% (interpolated values: P(Cluster)_70/95_ = 0.218, P(Epoch)_70/95_ = 0.046).

### Sensor-level waves

Next, we tested the sensitivity of our algorithm in detecting transient traveling wave events embedded in noise. For this, we first simulated physiological aperiodic activity following the same procedure as before. To this, we added (in the time domain, at the sensor level) the wave template as defined from the spherical phase gradient used in the fitting step of our algorithm (Figure 3A; for details, see Methods, *Simulation of EEG data: Sensor-level waves*). The rationale of this procedure is thus to recover the injected wave pattern under varying amounts of noise. Specifically, we varied the relative contributions of the source-level noise and white noise that was independently added to the signal of each electrode.

**Figure 3.**
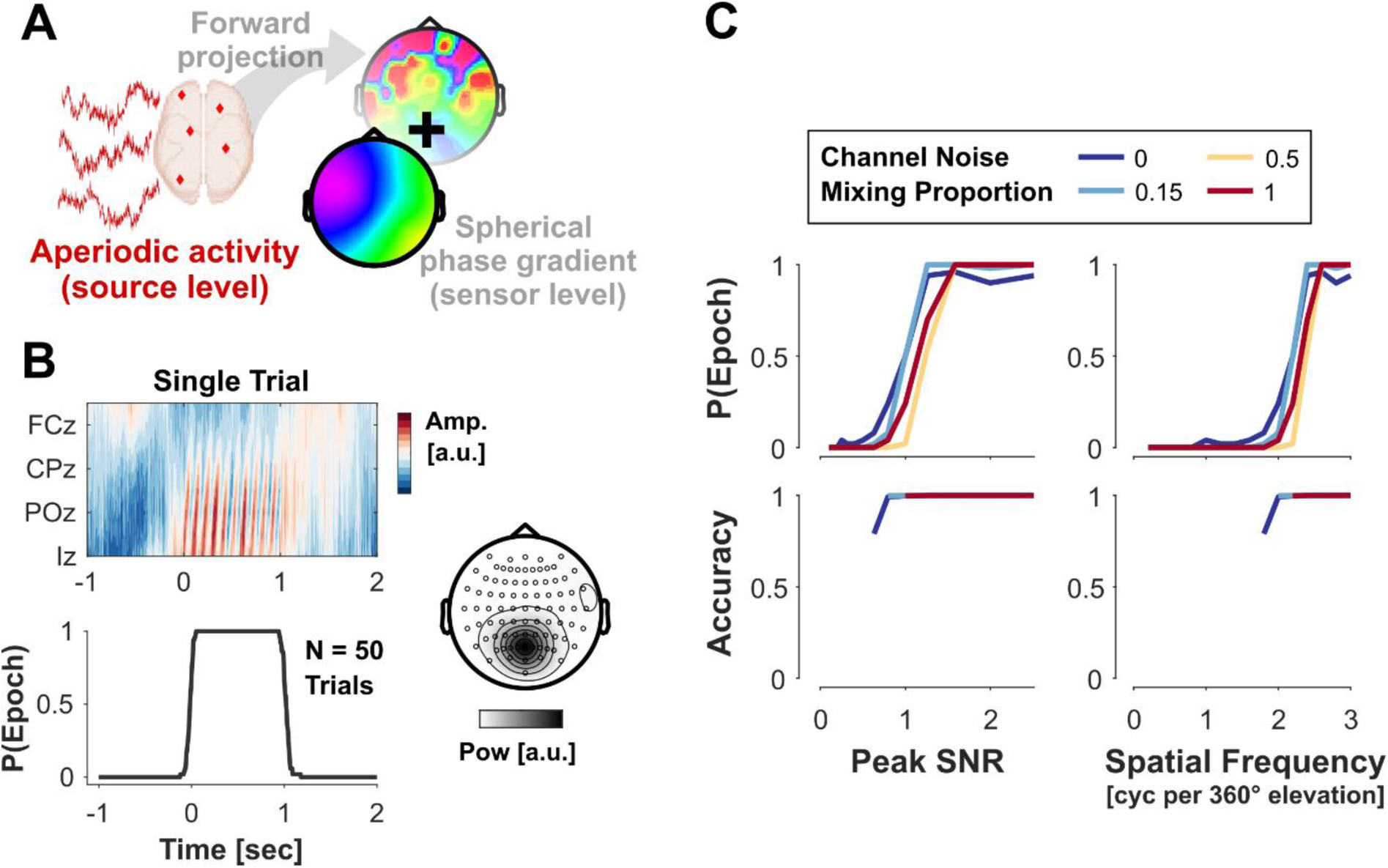
Validation of the wave detection algorithm with sensor-level simulated waves. A: Simulation procedure. Signals were simulated as the sum of forward-projected source-level aperiodic activity and a spherical wave template (localized in time-/frequency-/electrode-space). B: Top panel: simulated signal for a single trial, showing a 10 Hz forward wave pattern over occipital sensors between t = 0 and t = 1 sec. Bottom panel: detection probability of the same template across 50 simulated trials. Right: topography of mean oscillatory power (at 10 Hz) showing the spatial localization of the wave. C: Detection probabilities and accuracy of reconstructed wave direction for different parameter values. Metrics for each combination were computed over a set of 50 simulated trials, each comprising a 1-second wave event with a random wave direction.

The top panel of Figure 3B shows an example of a single trial with a 1-second simulated forward wave (posterior sensors leading in phase) localized over occipital regions. When evaluated over a set of simulated trials, the algorithm consistently detects a wave epoch in all cases, accurately recovering the time course of the ground-truth event (bottom panel).

For each set of tested parameters, we extracted the wave detection probability and the accuracy of the recovered wave direction. The results (summarized in Figure 3C) show that performance plateaus at high scores for both detection probability and accuracy above certain thresholds for the wave’s SNR and its spatial frequency.

These thresholds are only marginally affected by the type of noise, with slightly worse performance when the entire noise stems from the electrode. This may be explained by the fact that SNR here is based on the RMS of the signals, which is more strongly dominated by frequencies lower than the wave in the aperiodic activity as compared to the white noise.

Overall, these results demonstrate that the algorithm can reliably recover idealized wave patterns injected at the sensor level under variable noise conditions.

### Testing different configurations of source-level activity generating EEG alpha waves

After establishing that our algorithm can detect idealized waves defined at the sensor level, we now turn to more physiological cases in which the sensor-level wave pattern is generated by activity at the source level. Specifically, we follow previous studies (Schwenk & Alamia, 2025; Zhigalov & Jensen, 2023) that show planar traveling waves in the EEG can be explained by few oscillating dipoles with a consistent phase offset (illustrated in Figure 4A). To investigate which configurations of sources will lead to a wave being detected by our algorithm, we tested four different pathways. In all cases, we aimed to model the propagation of alpha rhythmic activity (temporal frequency F = 10Hz) in the visual hierarchy, i.e., along the dorsal (occipital-parietal) or ventral (occipital-temporal) visual pathway. The source locations forming the grid points of these pathways are shown in Figure 4B.

**Figure 4.**
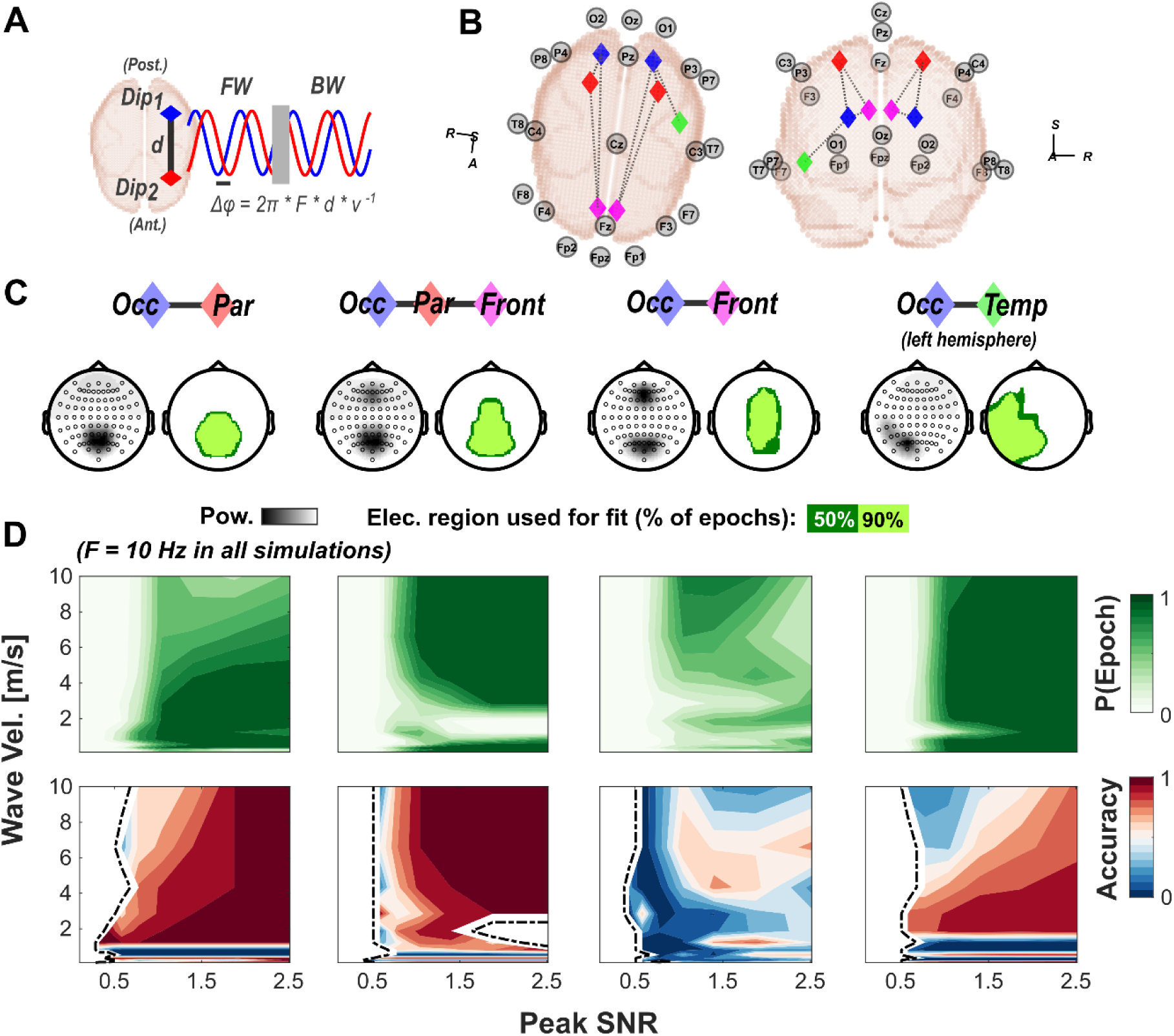
Results of the wave detection algorithm applied to forward simulations of phase-offset oscillating dipoles. A: Illustration of the studied wave-generating mechanism. A small number of dipoles (here: two) oscillate with a consistent phase offset that is either forward or backward directed (with respect to the visual hierarchy). The phase difference Δφ between dipoles is a function of the temporal frequency F (here, F = 10Hz), the physical distance between sources d, and the wave velocity v. B: Source locations underlying the four pathways tested below. Dashed lines illustrate all included direct connections between sources. Wherever sources are bilateral, left- and right-hemispherical activity was identical for each area. Positions of a subset of EEG sensors are given for reference; all simulations were evaluated on a full 64-channel EEG. Legend: A – Anterior; S – Superior; R – Right. C: Overview of the different pathways tested. Each subpanel shows topographical plots of the power distribution (at the wave frequency) and electrode regions where waves were detected (evaluated for SNR = 2, v = 5 m/s). D: Wave detection performance for each pathway (order of columns corresponding to panel C) over a grid of values for wave SNR and propagation velocity. Accuracy scores are defined as the proportion of waves correctly classified as FW/BW.

We simulated waves traveling along each pathway in either of the two possible directions, denoting as forward (FW) and backward (BW) the cases when lower-level areas are leading and trailing in phase, respectively. As for the sensor-level analysis, we evaluated performance for every combination of SNR and wave velocity in terms of detection probability and accuracy of the estimated wave direction (now classified as binary: FW/BW). Figure 4C & D summarizes the results of these simulations. For each pathway, our algorithm detected planar wave patterns over topographical regions with the highest oscillatory power (panel C). Notably, the two pathways involving frontal regions (*Occ-Par-Front* and *Occ-Front*) exhibit a bimodal power topography. In these cases, the two peaks are merged into a single cluster (either because the lower threshold for the spatial extent of the cluster connects them or through merging of clusters that coincide in time and frequency, see *Methods: Extraction of oscillatory clusters*), allowing for a long-range wave to be detected. However, as discussed further below, the parametrization of this type of spatiotemporal pattern as a wave is redundant, since the (directed) phase-offset between the two peaks can be extracted directly at the level of individual sensors. We still include these cases here to illustrate the behavior of our algorithm for a range of possible source-level configurations.

The detection metrics over the full grid of parameters (panel D) show that stable quantification of the wave between pathways differs mainly in the range of propagation velocities. Specifically, for the case of two sources in close proximity (*Occ-Par*) with a resulting unimodal power-topography, wave detection fails if propagation velocity is high (approx. > 6 m/s) because the range of phase values in the sensor space becomes too low to be reliably distinguished from a standing wave. A wider spatial extent of sensors (*Occ-Par-Front* / *Occ-Front and Occ-Temp*) supports wave detection also at higher velocities.

Notably, the split topography in the two pathways with frontal involvement corrupts the wave detection at low propagation velocities, most prominently in the case of only two distant sources (*Occ-Front*). This effect is explained by the inclusion of intermediate sensors between the two peaks: the phase values at these sensors are dominated by noise (since the amplitude of the wave signal is low) and therefore become increasingly better approximated the closer the wave fit approaches the null model of a standing wave (uniform phase). This highlights why power topographies with two or more distinct component in the sensor space limit the use cases for this type of detection algorithm. Implications of this limitation, as well as potential solutions, are discussed further below (see *Discussion: Source-level correlates of waves observed at the scalp* level).

In summary, our algorithm also detects planar traveling wave patterns when they are generated by phase-offset oscillating dipoles at the source level. Within a range of plausible physiological conduction delays (Girard et al., 2001; Lemaréchal et al., 2021; Schmolesky et al., 1998), the two directions on the main axis of propagation (FW/BW) can be reliably distinguished.

### Analysis of real EEG data

In the final step, we evaluated our algorithm using real human EEG data, specifically targeting alpha wave propagation in the visual system. First, we re-analyzed data from a previous study (Pang (庞兆阳) et al., 2020) that compared alpha traveling waves during stimulus on- and off-periods. For this dataset, alpha wave patterns have been reported based on two different quantification methods (2DFFT: (Pang (庞兆阳) et al., 2020), ROI-based approximative planar phase fitting: (Schwenk & Alamia, 2025)), making it a suitable confirmatory analysis to evaluate our new algorithm. Secondly, we studied the distributions of alpha wave direction during the resting state in two publicly available datasets (Babayan et al., 2019; Chenot, Hamery, Truninger, De Boissezon, et al., 2024).

The results of our analysis for the Pang et al. dataset are summarized in Figure 5A and B. The stimulation sequence for this experiment consisted of an alternation between 5 seconds of stimulation (a randomly flickering disk for the experimental condition included here) and 5 seconds of rest (a blank screen), while subjects were instructed to maintain fixation throughout. The time-frequency plots in panel A show the trial-average pattern of detected wave epochs for three different subjects. Here, the contours mark time-frequency points at which epochs were detected at all (in at least 5% of trials), while color represents the mean direction of those wave epochs (white areas showed no consistent wave direction across trials). As reported in the original publication, we observed consistent traveling waves in the alpha band in all subjects. At the subject level, the modulation by the stimulus showed some variability, with some subjects exhibiting a clear reversal from BW to FW waves (example S13), while others remained in the BW state (S7). Notably, and in addition to previously published results, we found that some subjects exhibited a modulation consistent with a reversal, but with FW and/or BW directions tilted away from the anterior-posterior axis (S2).

**Figure 5.**
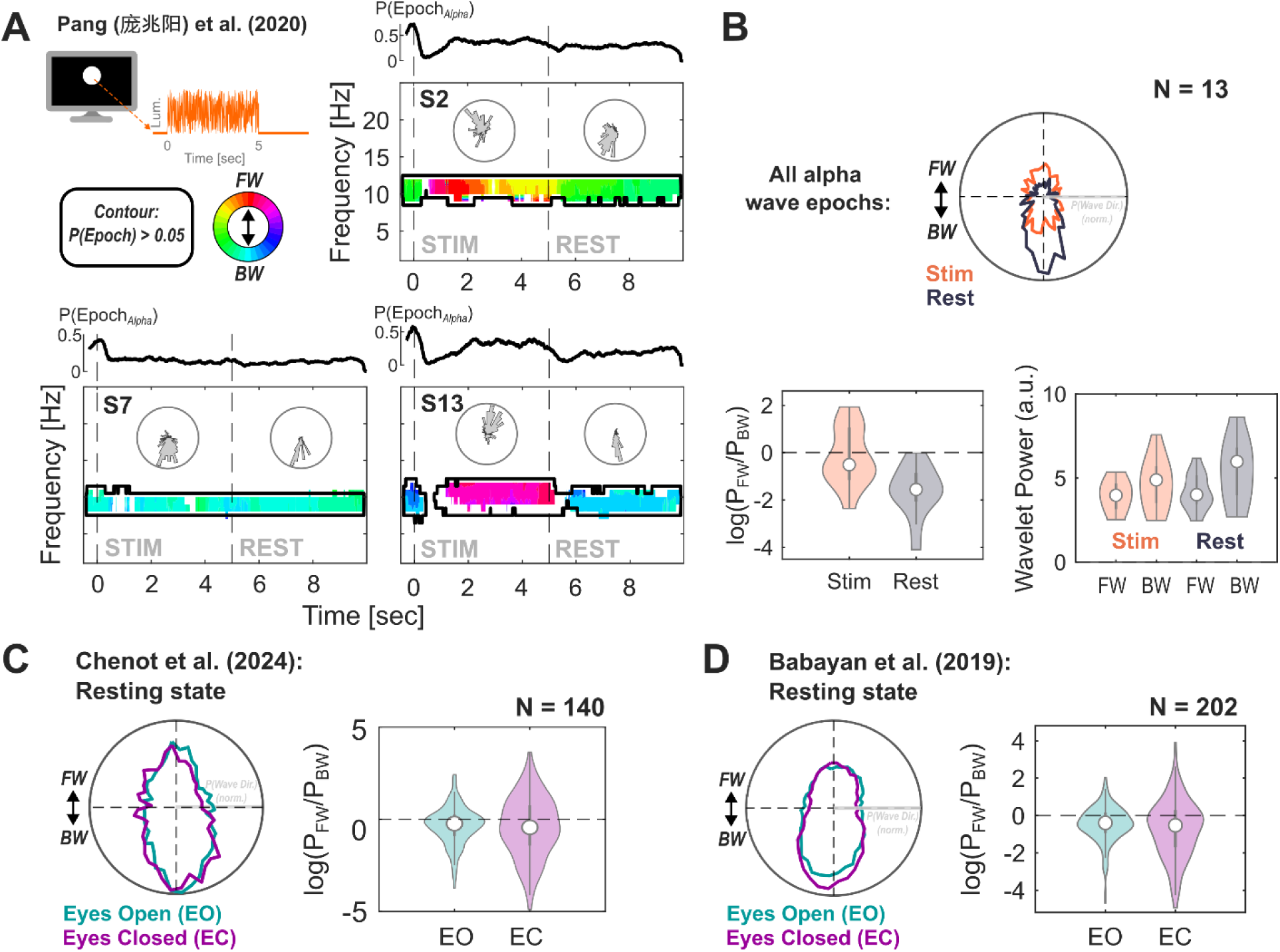
Results of the wave detection algorithm for three EEG datasets. A: Subject-level results for the data from Pang (庞兆阳) et al. (2020). Time-frequency plots display the detected epochs (averaged across trials) for three subjects, each highlighting a distinct pattern of modulation by the stimulus. Contours mark time-frequency regions with epochs detected in more than 5% of trials. Colored areas represent regions where the mean vector length of wave directions (between epochs) exceeded the random level (p < 0.05, uncorrected), with color indicating mean wave direction (see legend). Inset plots show histograms of wave direction for all alpha epochs within each time period. Small top panels show the probabilities of alpha epochs over the time-course of the trial. B: Population-level results for the Pang (庞兆阳) et al. (2020) dataset. Top: grand-average histograms of wave direction for each time window. Distributions are normalized as probabilities (per subject and time window) before averaging. Bottom, left: log-ratio of FW and BW wave-probabilities for the two time-windows. Values below zero indicate a bias towards the BW state. Right: Mean oscillatory power underlying individual wave epochs, separated by time window and wave direction. C, D: Results for alpha wave epochs in two publicly available datasets, separately for eyes-open (EO) and eyes-closed (EC) resting states. Left panels show the grand average histograms of wave direction, and right panels show the log-ratio of FW and BW wave-probabilities (as in B, top and bottom left panels).

The mean histogram of wave directions (Figure 5B, top, alpha-only) confirms that the rest-period was consistently BW, whereas both types of waves were present in the stimulus-period. We classified waves into the two directions (with a broad window of +/-45° to account for the observed off-axis deviations) and compared their log-ratio as a measure of bias towards either state between time-periods (Figure 5B, bottom left), which confirmed a significant modulation by the stimulation (Bayesian paired-samples t-test: BF_10_ = 111). Thus, our new method matches the results reported in the original publication (Pang (庞兆阳) et al., 2020), while offering much greater resolution in all three relevant dimensions (time-/frequency- and electrode-space).

An additional advantage of our algorithm over previous approaches, such as the 2D FFT method, is that it allows for the analysis of waves at level of single events (epochs). As an exploratory analysis, we compared oscillatory power between the two types of waves (FW/BW) and the two time-periods (Figure 5B, bottom right). This revealed that power during BW waves was consistently higher than that during FW waves, with a slightly larger divergence during the resting state (Bayesian repeated measures ANOVA, with both factors, Direction and Time-Window, and their interaction: BF_10_ > 21).

We also analyzed alpha wave direction in two publicly available resting-state datasets to assess the consistency of our result of a strong BW bias at rest across datasets. The results for these analyses are summarized in Figure 5C and D. For both datasets, the overall rates of alpha wave epochs (across trials and time-points) were in a range between 10% and 25%, similar to those observed in the Pang et al. dataset (P(Epoch_Alpha_), Mean +/-SD: Pang (庞兆阳) et al.: 0.142 +/-0.062; Chenot et al.: 0.243 +/-0.042; Babayan et al.: 0.104 +/-0.019). Similarly, the distribution of wave direction was again bimodal, aligning with the FW/BW axis. Waves in both conditions for both datasets were overall significantly biased towards the BW state (Bayesian t-tests for difference from zero-mean, EC for Chenot et al., BF_10_ = 8.9, all other BF_10_ > 132). However, this bias was considerably smaller than for the Pang et al. data (mean log-ratios between -0.355 and -0.729, compared to -1.625 in the rest period for Pang et al.), possibly due to differences in the experimental design (visual stimulation compared to resting states). Notably, a large proportion of subjects also exhibited a strong bias towards the FW state. We found no strong evidence for a difference in the FW/BW ratio between the two resting-state conditions in either dataset (EO vs. EC, both BF_10_ < 2).

In summary, the above analyses demonstrate that our algorithm is well-suited for studying traveling waves in real EEG data, providing relevant parameters (such as wave direction) with good consistency at high temporal resolution.

## Discussion

We have demonstrated that our proposed method, which performs wave template fitting based on clusters of oscillatory activity, provides a flexible quantification of transient traveling wave events. The algorithm shows good performance under conditions of physiological noise, and accurately recovers phase templates at the sensor level. Moreover, we demonstrated that it can reliably classify the directionality of large-scale waves traveling along visual pathways, based on the simplest model of phase-offset oscillating dipoles at the source level. We also replicated and expanded findings from a previously studied dataset and found consistent results when applying our analysis to two additional resting-state datasets.

In the following sections, we will discuss the advantages and limitations of our algorithm in comparison to other methods, and briefly highlight the possible implications of our simulations for the study of the source-level correlates of EEG traveling waves.

### Comparison to other methods

Our method is conceptually based on the planar template fitting introduced by Zhang et al. for analyzing ECoG data (Zhang et al., 2018). The methodological advancement here lies in adapting this procedure to the EEG, crucially accounting for the spherical shape of the sensor space and the noise background typical in EEG recordings. We will therefore compare our method to others that aim to quantify waves specifically in the EEG. However, it is worth noting the great variety of wave-detection methods that have been established for invasive data (e.g., (Alexander et al., 2013; Bolt et al., 2022; Muller et al., 2014; Takagaki et al., 2011; Townsend & Gong, 2018; Zhang et al., 2018)). Some of these may be adaptable to the EEG, and more work generalizing across data sources is likely to be expected in the future.

Our method predefines a set of wave templates to which datapoints can be matched, i.e., it follows a model-based approach. As such, we primarily evaluate it in relation to other model-based methods. Further below, we briefly discuss the differences between this method and other data-driven methods.

A commonly used method to quantify waves in the EEG is the analysis of 2D-FFT spectra along a predefined line of electrodes (e.g., the midline) (Alamia et al., 2020, 2023, 2024; Alamia & VanRullen, 2019; Luo & Ester, 2025; Pang (庞兆阳) et al., 2020; Tarasi et al., 2025; Zeng et al., 2024). There, waves are quantified based on the proportion of energy within the quadrants of the 2D-FFT spectrum corresponding to the two possible directions on the line of electrodes. While simple in implementation, this method is limited to one axis of propagation at a time and offers poor spatial and temporal resolution. Our method notably improves on this in all aspects, allowing for the quantification of the wave direction in full 360°, at high temporal resolution in time-frequency space (as given by the wavelet transform). The resolution in spatial frequency is improved by the inclusion of more electrodes and sampling over two dimensions instead of just one. Another limitation of the 2D-FFT method is that its wave quantification uses oscillatory power. While a normalization based on data shuffling (introduced in (Pang (庞兆阳) et al., 2020)) can be used to correct for overall power fluctuations in the data, this correction is constrained by the same low resolution as the 2D-FFT itself and cannot entirely eliminate the confound between oscillatory power and extracted wave power. Our method, instead, pre-selects data based on power (clustering), while the wave estimate itself relies only on phase. This approach effectively confines the influence of power to the probability of detecting a wave (i.e., an epoch), whereas parameters of the extracted wave should be largely independent.

Analogous to the 2D-FFT approach, some studies have also employed phase-fitting methods on lines of electrodes, with a simple line fit as the underlying wave model (Burkitt et al., 2000; Ito et al., 2005; Patten et al., 2012). As discussed above, our method extends this approach to a 2D space of electrodes. Additionally, our wave template definition takes into account the spherical shape of the head. This eliminates the spherical distortion introduced when parts of the curved surface are approximated by a line or planar slopes (above studies and (Schwenk & Alamia, 2025)).

As mentioned above, data-driven methods provide an alternative approach to detecting traveling waves. Broadly speaking, these methods are all built on some decomposition of the spatial phase map (or gradient) into components or motifs (Alexander et al., 2006, 2013; Li et al., 2025; Patten et al., 2012; Petras et al., 2025). These components can then be matched to interpretable waveforms (such as the planar waves targeted by our method) *post hoc*. This makes such methods less biased than our method, allowing for a comprehensive view of all patterns present in the data (as opposed to only the one tested, in our approach). Conversely, they are not optimized to test hypotheses about the transient occurrence of specific wave patterns, particularly when the matching between extracted components and the targeted wave type is ambiguous, or when the latter does not occur frequently or consistently enough to be reliably extracted.

Notably, however, the results from our analysis for one resting-state dataset (Babayan et al., 2019) match those obtained by Li et al. for the same dataset using a data-driven approach (Weakly Orthogonal Conjugate Contrast Analysis (WOCCA); (Li et al., 2025)). The authors of that study also report a predominance of waves aligned with the anterior-posterior axis, with an overall bias towards the backward state that was significantly stronger in the eyes-closed (EC) condition (the comparison EC vs. EO in our analysis shows the same trend but no significant effect, cf. Figure 5D). This suggests that findings about traveling wave dynamics (at least for a dominant component such as resting-state alpha) generalize across different methods of quantification.

In summary, our algorithm outperforms previous model-based methods and can be viewed as complementary to data-driven methods, with a more targeted use case.

### Method limitations

A central limitation of our approach is that it targets only a single wave type, i.e., planar waves (in principle, the proposed framework is applicable to any wave that can be formulated as a template; however, we discuss here based only on our simulations). In addition to building on previous EEG findings (Alamia & VanRullen, 2019; Li et al., 2025; Lozano-Soldevilla & VanRullen, 2019; Petras et al., 2025), this focus is also theoretically motivated by the ubiquity of planar waves at the cortical level. Here, synchronized wavefronts with a single direction have been observed as large-scale spontaneous waves (Bahramisharif et al., 2013; Davis et al., 2020; Halgren et al., 2019; Zhang et al., 2018), and as movement-related events in the motor cortex (Balasubramanian et al., 2020; Best et al., 2017; Rubino et al., 2006; Takahashi et al., 2011, 2015). Similarly, stimulus evoked responses in sensory cortices have been shown to elicit radiating waves propagating outward from the first activation (Chemla et al., 2019; Muller et al., 2014, 2018; Sato et al., 2012), which would likely result in planar traveling waves in the EEG.

However, a variety of other wave types have also been characterized. Most prominent among these are rotational patterns, observed at the mesoscale (Alexander & Dugué, 2025; Das et al., 2024; Huang et al., 2010; Muller et al., 2016; Rule et al., 2018) and at scalp-level (Liu et al., 2018; Xu et al., 2025). More complex wave patterns forming around single critical points (phase singularities) can be described at the mesoscale (Townsend et al., 2015; Townsend & Gong, 2018), but their possible projection to the sensor space remains unclear.

Although our method does not specifically target these wave types, their potential occurrence in the EEG may impact its reliability. In theory, any pattern that does not match the underlying template should result in poor fits and not be classified. However, erroneous classification could occur in practice, in particular if the initially extracted cluster only covers parts of a larger pattern (for example, the lateral edges of a rotating pattern can be locally approximated by a planar wave). This type of interference may be counteracted by parallel detection of other wave types, or *post-hoc* rejection of extracted epochs based on additional criteria (e.g., detection of phase singularities in close proximity to the cluster region).

Another limitation of our method is that it requires a reasonably high SNR for the wave to be detected. This is because the extent of the electrode region (based on oscillatory power) that is matched with the template impacts the goodness of fit. Additionally, for clusters that are small in electrode space, the fit may not achieve a sufficient spread in phase angles, allowing it to be distinguished from a standing wave, even if the fit is good. Adjustment of the parameters (cluster thresholds, use of ROI and/or FOI) allows some control over the trade-off between sensitivity and specificity. However, the target use case for our algorithm is the analysis of strong oscillatory EEG components (e.g., alpha, theta). Other methods may be more suitable for studying wave dynamics in low-SNR signals, such as data-driven approaches (see *Discussion: Comparison to other methods*).

Lastly, since our method bases its phase estimate on the wavelet transform (WT), it is potentially susceptible to distortion introduced by non-sinusoidal waveforms. Previous studies have shown that some oscillatory components in the EEG deviate from the sinusoidal waveform, and that this constitutes a physiologically (and potentially functionally) meaningful feature (Cole & Voytek, 2017, 2019; Schaworonkow & Nikulin, 2019). In most cases, the wave detection in our algorithm should be robust against the non-sinusoidal nature of the underlying oscillation. The phase estimate is extracted from the inverse WT transform of the entire cluster (see *General description of the* method*: Extraction of oscillatory clusters*), which allows for small irregularities but will be dominated by the base oscillation. However, the WT of a regularly repeating non-sinusoidal waveform typically splits into harmonic components of the base frequency. Our algorithm may extract these components as separate clusters and subsequently classify them as simultaneous waves in different frequency bands (given that both are in the analyzed frequency range and each follows a consistent phase pattern). As with any Fourier- or wavelet-based analysis, careful inspection of the data is therefore warranted if separate wave components are present at integer-multiple frequencies.

### Source-level correlates of waves observed at the scalp level

The focus of the present work was on establishing a reliable wave detection at the sensor level. Since our method does not include a source reconstruction step, any phase pattern matching a planar wave will be classified as an epoch by our algorithm, regardless of its generating mechanism at the source level.

Direct inference about the source-level activity underlying observed scalp-level waves is problematic, and previous studies attempting this have relied on various forms of wave models based on physiological assumptions (Grabot et al., 2025; Hindriks et al., 2014; Lozano-Soldevilla & VanRullen, 2019; Petras et al., 2025; Zhigalov & Jensen, 2023). Similarly, here and in previous work (Alamia & VanRullen, 2019; Schwenk & Alamia, 2025), we have explored the scalp-level projections of various configurations of phase-offset oscillating dipoles. The underlying generator model simulates waves traveling down- or upstream along the visual hierarchy. Our results from those simulations showed that planar wave patterns could arise from all tested pathways (comprising occipital, parietal, frontal and temporal sources), confirming our own and others’ findings that a small number of oscillating sources is the most parsimonious explanation for this type of wave pattern ((Orsher et al., 2024; Schwenk & Alamia, 2025; Zhigalov & Jensen, 2023), but see (Hindriks et al., 2014) for a different account).

Our results further showed that configurations with sources at close physical distance require slower propagation velocities for the wave to be reliably detected at the scalp. For the two pathways with contiguous topographical distributions on the scalp (*Occ-Par* and *Occ-Temp*), the parameter range of detected waves matches the range of inter-areal conduction velocities found in macaque and human cortex (Girard et al., 2001; Lemaréchal et al., 2021; Schmolesky et al., 1998). It should be noted, however, that our generator model assumes straight connections between areas. Additionally, neural integration times and local processing at each area are not taken into account. Both lead to an underestimation of the required axonal conduction velocities, rendering waves at the upper end of our tested parameter range less likely to occur physiologically (assuming direct cortico-cortical propagation). Nonetheless, our results suggest that topographically small oscillatory clusters may appear as a standing wave even if a (fast) traveling wave is present at the source level.

On the other extreme of the tested cases, we found that our algorithm can theoretically classify propagation direction for source configurations that project to distinct topographical clusters on the scalp (*Occ-Par-Front* and *Occ-Front*), although this was not reliable for more physiologically plausible propagation velocities. As indicated above, the case of topographically split clusters (each with a uniform phase) is, of course, more succinctly quantified by the phase offset between them. Additional steps applied prior to phase fitting (e.g., splitting of bimodal clusters) may be used to control the processing of these cases in the analysis pipeline. However, in analysis contexts where waves can be assumed to be generated strictly through cortico-cortical propagation, it may be necessary to generalize between cases of contiguous and bimodal (split) scalp topographies. Thus, ultimately, the quantification also depends on the definition of a wave that is applied.

While our simulations relied on the cortico-cortical propagation model, this is not the only account for waves observed in the EEG. Other studies have shown that mesoscale waves, i.e., those traveling along horizontal connections within one cortical area, can lead to similar wave patterns at the scalp level (Grabot et al., 2025; Hindriks et al., 2014; Petras et al., 2025). Interestingly, in the study by Petras et al., the modulation of the wave component associated with meso-scale waves (which they induced visually) was limited to the stimulation frequency, while alpha waves were not modulated. Assuming a long-range cortico-cortical propagation mechanism for alpha (Alamia & VanRullen, 2019; Schwenk & Alamia, 2025; Zhigalov & Jensen, 2023), this suggests that the two types of waves (meso-scale and cortico-cortical) may be separable in the EEG. The wave parametrization in our algorithm would be well-suited for this type of analysis. Specifically, different wave scales may be separable into different spatial frequency bands. Furthermore, by utilizing the high temporal resolution, our analysis would enable a correlation between wave direction and a variable wave-inducing stimulus.

### Conclusions

We presented an analysis pipeline for detecting and quantifying planar traveling waves in the EEG, based on a combination of oscillatory clustering and a spherical wave fit. Our validation tests and forward simulations showed that our algorithm performs well under physiologically realistic conditions. We further discussed the limitations of our method’s use cases in the context of different possible source-level mechanisms that generate waves in the EEG. In comparison with other approaches, we demonstrated that our method outperforms previous model-based analyses by allowing for more flexible wave quantification across all three data dimensions (time/frequency/electrode space).

## Code and data availability

The code used to generate the results presented in this study is publicly available (https://github.com/jcbschwenk/eeg-planarwave-clust). Any additional information is available from the corresponding author upon request.

## Acknowledgments

This project was funded by the European Union under the European Union’s Horizon 2020 research and innovation program (grant agreements No. 101075930 to AA). The funders had no role in study design, data collection and analysis, decision to publish, or preparation of the manuscript. Views and opinions expressed are those of the author(s) only and do not necessarily reflect those of the European Union or the European Research Council (ERC). Neither the European Union nor the granting authority can be held responsible for them. The authors are grateful to Martina Pasqualetti for helpful feedback on the analysis code, as well as Leslie Marie-Louise for administrative support.

## References

Alamia, A., Gordillo, D., Chkonia, E., Roinishvili, M., Cappe, C., & Herzog, M. H. (2024). Oscillatory Traveling Waves Provide Evidence for Predictive Coding Abnormalities in Schizophrenia. Biological Psychiatry, S0006322324017827. 10.1016/j.biopsych.2024.11.014

Alamia, A., Terral, L., D’ambra, M. R., & VanRullen, R. (2023). Distinct roles of forward and backward alpha-band waves in spatial visual attention. eLife, 12, e85035. 10.7554/eLife.85035

Alamia, A., Timmermann, C., Nutt, D. J., VanRullen, R., & Carhart-Harris, R. L. (2020). DMT alters cortical travelling waves. eLife, 9, e59784. 10.7554/eLife.59784

Alamia, A., & VanRullen, R. (2019). Alpha oscillations and traveling waves: Signatures of predictive coding? PLOS Biology, 17(10), e3000487. 10.1371/journal.pbio.3000487

Alexander, D. M., & Dugué, L. (2025). The dominance of large-scale phase dynamics in human cortex, from delta to gamma (p. 2024.06.04.597334). bioRxiv. 10.1101/2024.06.04.597334

Alexander, D. M., Jurica, P., Trengove, C., Nikolaev, A. R., Gepshtein, S., Zvyagintsev, M., Mathiak, K., Schulze-Bonhage, A., Ruescher, J., Ball, T., & van Leeuwen, C. (2013). Traveling waves and trial averaging: The nature of single-trial and averaged brain responses in large-scale cortical signals. NeuroImage, 73, 95–112. 10.1016/j.neuroimage.2013.01.016

Alexander, D. M., Trengove, C., Wright, J. J., Boord, P. R., & Gordon, E. (2006). Measurement of phase gradients in the EEG. Journal of Neuroscience Methods, 156(1), 111–128. 10.1016/j.jneumeth.2006.02.016

Babayan, A., Erbey, M., Kumral, D., Reinelt, J. D., Reiter, A. M. F., Röbbig, J., Schaare, H. L., Uhlig, M., Anwander, A., Bazin, P.-L., Horstmann, A., Lampe, L., Nikulin, V. V., Okon-Singer, H., Preusser, S., Pampel, A., Rohr, C. S., Sacher, J., Thöne-Otto, A., … Villringer, A. (2019). A mind-brain-body dataset of MRI, EEG, cognition, emotion, and peripheral physiology in young and old adults. Scientific Data, 6(1), 180308. 10.1038/sdata.2018.308

Bahramisharif, A., Gerven, M. A. J. van, Aarnoutse, E. J., Mercier, M. R., Schwartz, T. H., Foxe, J. J., Ramsey, N. F., & Jensen, O. (2013). Propagating Neocortical Gamma Bursts Are Coordinated by Traveling Alpha Waves. Journal of Neuroscience, 33(48), 18849–18854. 10.1523/JNEUROSCI.2455-13.2013

Balasubramanian, K., Papadourakis, V., Liang, W., Takahashi, K., Best, M. D., Suminski, A. J., & Hatsopoulos, N. G. (2020). Propagating Motor Cortical Dynamics Facilitate Movement Initiation. Neuron, 106(3), 526–536.e4. 10.1016/j.neuron.2020.02.011

Best, M. D., Suminski, A. J., Takahashi, K., Brown, K. A., & Hatsopoulos, N. G. (2017). Spatio-Temporal Patterning in Primary Motor Cortex at Movement Onset. Cerebral Cortex, 27(2), bhv327. 10.1093/cercor/bhv327

Bolt, T., Nomi, J. S., Bzdok, D., Salas, J. A., Chang, C., Thomas Yeo, B. T., Uddin, L. Q., & Keilholz, S. D. (2022). A parsimonious description of global functional brain organization in three spatiotemporal patterns. Nature Neuroscience, 25(8), 1093–1103. 10.1038/s41593-022-01118-1

Burkitt, G. R., Silberstein, R. B., Cadusch, P. J., & Wood, A. W. (2000). Steady-state visual evoked potentials and travelling waves. Clinical Neurophysiology, 111(2), 246–258. 10.1016/S1388-2457(99)00194-7

Chemla, S., Reynaud, A., di Volo, M., Zerlaut, Y., Perrinet, L., Destexhe, A., & Chavane, F. (2019). Suppressive traveling waves shape representations of illusory motion in primary visual cortex of awake primate. The Journal of Neuroscience, 39(22), 2792–18. 10.1523/JNEUROSCI.2792-18.2019

Chenot, Q., Hamery, C., Truninger, M., De Boissezon, X., Langer, N., & Scannella, S. (2024). EEG Resting-state Microstates Correlates of Executive Functions [Dataset]. Openneuro. 10.18112/OPENNEURO.DS005305.V1.0.1

Chenot, Q., Hamery, C., Truninger, M., Langer, N., De boissezon, X., & Scannella, S. (2024). Investigating the relationship between resting-state EEG microstates and executive functions: A null finding. Cortex, 178, 1–17. 10.1016/j.cortex.2024.05.019

Cole, S. R., & Voytek, B. (2017). Brain Oscillations and the Importance of Waveform Shape. Trends in Cognitive Sciences, 21(2), 137–149. 10.1016/j.tics.2016.12.008

Cole, S. R., & Voytek, B. (2019). Cycle-by-cycle analysis of neural oscillations. Journal of Neurophysiology, jn.00273.2019. 10.1152/jn.00273.2019

Das, A., Zabeh, E., Ermentrout, B., & Jacobs, J. (2024). Planar, Spiral, and Concentric Traveling Waves Distinguish Cognitive States in Human Memory (p. 2024.01.26.577456). bioRxiv. 10.1101/2024.01.26.577456

Davis, Z. W., Muller, L., Martinez-Trujillo, J., Sejnowski, T., & Reynolds, J. H. (2020). Spontaneous travelling cortical waves gate perception in behaving primates. Nature, 587(7834), 432–436. 10.1038/s41586-020-2802-y

Delorme, A., & Makeig, S. (2004). EEGLAB: An open source toolbox for analysis of single-trial EEG dynamics including independent component analysis. Journal of Neuroscience Methods, 134(1), 9–21. 10.1016/j.jneumeth.2003.10.009

Donoghue, T., Haller, M., Peterson, E. J., Varma, P., Sebastian, P., Gao, R., Noto, T., Lara, A. H., Wallis, J. D., Knight, R. T., Shestyuk, A., & Voytek, B. (2020). Parameterizing neural power spectra into periodic and aperiodic components. Nature Neuroscience, 23(12), 1655–1665. 10.1038/s41593-020-00744-x

Fakche, C., & Dugué, L. (2024). Perceptual Cycles Travel Across Retinotopic Space. Journal of Cognitive Neuroscience, 36(1), 200–216. 10.1162/jocn_a_02075

Fellinger, R., Gruber, W., Zauner, A., Freunberger, R., & Klimesch, W. (2012). Evoked traveling alpha waves predict visual-semantic categorization-speed. NeuroImage, 59(4), 3379–3388. 10.1016/j.neuroimage.2011.11.010

Girard, P., Hupé, J. M., & Bullier, J. (2001). Feedforward and Feedback Connections Between Areas V1 and V2 of the Monkey Have Similar Rapid Conduction Velocities. Journal of Neurophysiology, 85(3), 1328–1331. 10.1152/jn.2001.85.3.1328

Grabot, L., Merholz, G., Winawer, J., Heeger, D. J., & Dugué, L. (2025). Traveling waves in the human visual cortex: An MEG-EEG model-based approach. PLOS Computational Biology, 21(4), e1013007. 10.1371/journal.pcbi.1013007

Halgren, M., Ulbert, I., Bastuji, H., Fabó, D., Erőss, L., Rey, M., Devinsky, O., Doyle, W. K., Mak-McCully, R., Halgren, E., Wittner, L., Chauvel, P., Heit, G., Eskandar, E., Mandell, A., & Cash, S. S. (2019). The generation and propagation of the human alpha rhythm. Proceedings of the National Academy of Sciences, 116(47), 23772–23782. 10.1073/pnas.1913092116

Hindriks, R., van Putten, M. J. A. M., & Deco, G. (2014). Intra-cortical propagation of EEG alpha oscillations. NeuroImage, 103, 444–453. 10.1016/j.neuroimage.2014.08.027

Huang, X., Xu, W., Liang, J., Takagaki, K., Gao, X., & Wu, J. (2010). Spiral Wave Dynamics in Neocortex. Neuron, 68(5), 978–990. 10.1016/j.neuron.2010.11.007

Ito, J., Nikolaev, A. R., & Leeuwen, C. van. (2005). Spatial and temporal structure of phase synchronization of spontaneous alpha EEG activity. Biological Cybernetics, 92(1), 54–60. 10.1007/s00422-004-0533-z

Lemaréchal, J.-D., Jedynak, M., Trebaul, L., Boyer, A., Tadel, F., Bhattacharjee, M., Deman, P., Tuyisenge, V., Ayoubian, L., Hugues, E., Chanteloup-Forêt, B., Saubat, C., Zouglech, R., Reyes Mejia, G. C., Tourbier, S., Hagmann, P., Adam, C., Barba, C., Bartolomei, F., … David, O. (2021). A brain atlas of axonal and synaptic delays based on modelling of cortico-cortical evoked potentials. Brain, 145(5), 1653–1667. 10.1093/brain/awab362

Li, Y., Qu, J., Yi, D., & Hong, B. (2025). Resolving Whole-Brain Alpha Traveling Waves Across Brain States. NeuroImage, 121538. 10.1016/j.neuroimage.2025.121538

Liu, X., Lu, Y., & Kuzum, D. (2018). High-Density Porous Graphene Arrays Enable Detection and Analysis of Propagating Cortical Waves and Spirals. Scientific Reports, 8(1), 17089. 10.1038/s41598-018-35613-y

Lozano-Soldevilla, D., & VanRullen, R. (2019). The Hidden Spatial Dimension of Alpha: 10-Hz Perceptual Echoes Propagate as Periodic Traveling Waves in the Human Brain Report The Hidden Spatial Dimension of Alpha: 10-Hz Perceptual Echoes Propagate as Periodic Traveling Waves in the Human Brain. CellReports, 26(2), 374–380.e4. 10.1016/j.celrep.2018.12.058

Luo, C., & Ester, E. F. (2025). Traveling waves link human visual and frontal cortex during working memory–guided behavior. Proceedings of the National Academy of Sciences, 122(30), e2415573122. 10.1073/pnas.2415573122

Muller, L., Chavane, F., Reynolds, J., & Sejnowski, T. J. (2018). Cortical travelling waves: Mechanisms and computational principles. Nature Reviews Neuroscience, 19(5), 255–268. 10.1038/nrn.2018.20

Muller, L., Piantoni, G., Koller, D., Cash, S. S., Halgren, E., & Sejnowski, T. J. (2016). Rotating waves during human sleep spindles organize global patterns of activity that repeat precisely through the night. eLife, 5, e17267. 10.7554/eLife.17267

Muller, L., Reynaud, A., Chavane, F., & Destexhe, A. (2014). The stimulus-evoked population response in visual cortex of awake monkey is a propagating wave. Nature Communications, 5. 10.1038/ncomms4675

Oostenveld, R., Fries, P., Maris, E., & Schoffelen, J. M. (2011). FieldTrip: Open source software for advanced analysis of MEG, EEG, and invasive electrophysiological data. Computational Intelligence and Neuroscience, 2011. 10.1155/2011/156869

Orsher, Y., Rom, A., Perel, R., Lahini, Y., Blinder, P., & Shein-Idelson, M. (2024). Sequentially activated discrete modules appear as traveling waves in neuronal measurements with limited spatiotemporal sampling. eLife, 12, RP92254. 10.7554/eLife.92254

Pang (庞兆阳), Z., Alamia, A., & VanRullen, R. (2020). Turning the Stimulus On and Off Changes the Direction of α Traveling Waves. Eneuro, 7(6), ENEURO.0218-20.2020. 10.1523/ENEURO.0218-20.2020

Patten, T. M., Rennie, C. J., Robinson, P. A., & Gong, P. (2012). Human Cortical Traveling Waves: Dynamical Properties and Correlations with Responses. PLOS ONE, 7(6), e38392. 10.1371/journal.pone.0038392

Petras, K., Grabot, L., & Dugué, L. (2025). Locally Induced Traveling Waves Generate Globally Observable Traveling Waves. Journal of Neuroscience, 45(34). 10.1523/JNEUROSCI.0089-25.2025

Rubino, D., Robbins, K. A., & Hatsopoulos, N. G. (2006). Propagating waves mediate information transfer in the motor cortex. Nature Neuroscience, 9(12), 1549– 1557. 10.1038/nn1802

Rule, M. E., Vargas-Irwin, C., Donoghue, J. P., & Truccolo, W. (2018). Phase reorganization leads to transient β-LFP spatial wave patterns in motor cortex during steady-state movement preparation. Journal of Neurophysiology, 119(6), 2212–2228. 10.1152/jn.00525.2017

Sato, T. K., Nauhaus, I., & Carandini, M. (2012). Traveling Waves in Visual Cortex. Neuron, 75(2), 218–229. 10.1016/j.neuron.2012.06.029

Schaworonkow, N., & Nikulin, V. V. (2019). Spatial neuronal synchronization and the waveform of oscillations: Implications for EEG and MEG. PLOS Computational Biology, 15(5), e1007055. 10.1371/journal.pcbi.1007055

Schmolesky, M. T., Wang, Y., Hanes, D. P., Thompson, K. G., Leutgeb, S., Schall, J. D., & Leventhal, A. G. (1998). Signal Timing Across the Macaque Visual System. Journal of Neurophysiology, 79(6), 3272–3278. 10.1152/jn.1998.79.6.3272

Schwenk, J. C. B., & Alamia, A. (2025). A hierarchical multiscale model of forward and backward alpha-band traveling waves in the visual system. PLOS Computational Biology, 21(8), e1013294. 10.1371/journal.pcbi.1013294

Takagaki, K., Zhang, C., Wu, J.-Y., & Ohl, F. W. (2011). Flow detection of propagating waves with temporospatial correlation of activity. Journal of Neuroscience Methods, 200(2), 207–218. 10.1016/j.jneumeth.2011.05.023

Takahashi, K., Kim, S., Coleman, T. P., Brown, K. A., Suminski, A. J., Best, M. D., & Hatsopoulos, N. G. (2015). Large-scale spatiotemporal spike patterning consistent with wave propagation in motor cortex. Nature Communications, 6(1), 7169. 10.1038/ncomms8169

Takahashi, K., Saleh, M., Penn, R. D., & Hatsopoulos, N. G. (2011). Propagating Waves in Human Motor Cortex. Frontiers in Human Neuroscience, 5, 40. 10.3389/fnhum.2011.00040

Tarasi, L., Alamia, A., & Romei, V. (2025). Perceptual Bias in Motion Discrimination is Related to Asymmetric Interhemispheric Alpha Traveling Waves. Advanced Science, e14623. 10.1002/advs.202414623

Townsend, R. G., & Gong, P. (2018). Detection and analysis of spatiotemporal patterns in brain activity. PLOS Computational Biology, 14(12), e1006643. 10.1371/journal.pcbi.1006643

Townsend, R. G., Solomon, S. S., Chen, S. C., Pietersen, A. N. J., Martin, P. R., Solomon, S. G., & Gong, P. (2015). Emergence of Complex Wave Patterns in Primate Cerebral Cortex. Journal of Neuroscience, 35(11), 4657–4662. 10.1523/jneurosci.4509-14.2015

Tzourio-Mazoyer, N., Landeau, B., Papathanassiou, D., Crivello, F., Etard, O., Delcroix, N., Mazoyer, B., & Joliot, M. (2002). Automated Anatomical Labeling of Activations in SPM Using a Macroscopic Anatomical Parcellation of the MNI MRI Single-Subject Brain. NeuroImage, 15(1), 273–289. 10.1006/nimg.2001.0978

Xu, Y., McInnes, A., Kao, C.-H., D’Rozario, A., Feng, J., & Gong, P. (2025). Spatiotemporal dynamics of sleep spindles form spiral waves that predict overnight memory consolidation and age-related memory decline. Communications Biology, 8(1), 1014. 10.1038/s42003-025-08447-4

Zeng, Y., Sauseng, P., & Alamia, A. (2024). Alpha Traveling Waves during Working Memory: Disentangling Bottom-Up Gating and Top-Down Gain Control. Journal of Neuroscience, 44(50). 10.1523/JNEUROSCI.0532-24.2024

Zhang, H., Watrous, A. J., Patel, A., & Jacobs, J. (2018). Theta and Alpha Oscillations Are Traveling Waves in the Human Neocortex. Neuron, 98(6), 1269–1281.e4. 10.1016/j.neuron.2018.05.019

Zhigalov, A., & Jensen, O. (2023). Perceptual echoes as travelling waves may arise from two discrete neuronal sources. NeuroImage, 272, 120047. 10.1016/j.neuroimage.2023.120047

